# Activity-dependent survival of odorant receptor neurons in ants

**DOI:** 10.1101/2023.10.04.560961

**Authors:** Bogdan Sieriebriennikov, Kayli R Sieber, Olena Kolumba, Jakub Mlejnek, Shadi Jafari, Hua Yan

## Abstract

Olfaction is essential for complex social behavior in eusocial insects. To discriminate complex social cues, ants evolved an expanded number of *odorant receptor* (*Or*) genes. Unlike most insect species, mutations in the obligate odorant co-receptor gene *orco* lead to loss of ∼80% antennal lobe glomeruli in ants. However, its cellular mechanism remains unclear. Here we demonstrate that this surprising neuronal phenotype results from massive apoptosis of odorant receptor neurons (ORNs) in the mid- to late-stages of pupal development. Further bulk and single-nucleus transcriptome analysis show that, although the majority of *orco*-expressing ORNs die in *orco* mutants, a small proportion of them survive: they express *ionotropic receptor* (*Ir*) genes that form IR complexes. In addition, we found that some *Or* genes are expressed in mechanosensory neurons as well as non-neuronal cells, possibly due to the leaky regulation from nearby non-*Or* genes. Our findings suggest that chemosensory receptors are required for activity-dependent survival of developing ORNs in ants.

## INTRODUCTION

The emergence of eusociality is a major evolutionary transition (*1*). Eusociality refers to a complex social system in which overlapping generations live together, wherein one or a few colony members reproduce and others cooperate for brood care or other specialized tasks, known as division of labor (*2*). This system is observed in several insect families, including ants, bees, and wasps in the order Hymenoptera, as well as termites, which have all evolved highly cooperative social behavior. In these insects, social recognition is critical for maintaining proper colony function. Sensory systems, and chemosensory communication via pheromones in particular, play a crucial role in social behavior by allowing individuals to recognize and communicate with each other. Antennae are the major anatomical organs involved in insect chemosensation. They possess hair-like structures called sensilla, which house the dendrites of odorant receptor neurons (ORNs) (Fig. 1A). The pheromone-sensing role of ORNs is essential for eusocial insects to achieve social recognition. Thus, deciphering the mechanisms underlying social communication requires understanding the development and function of ORNs.

**Figure 1.**
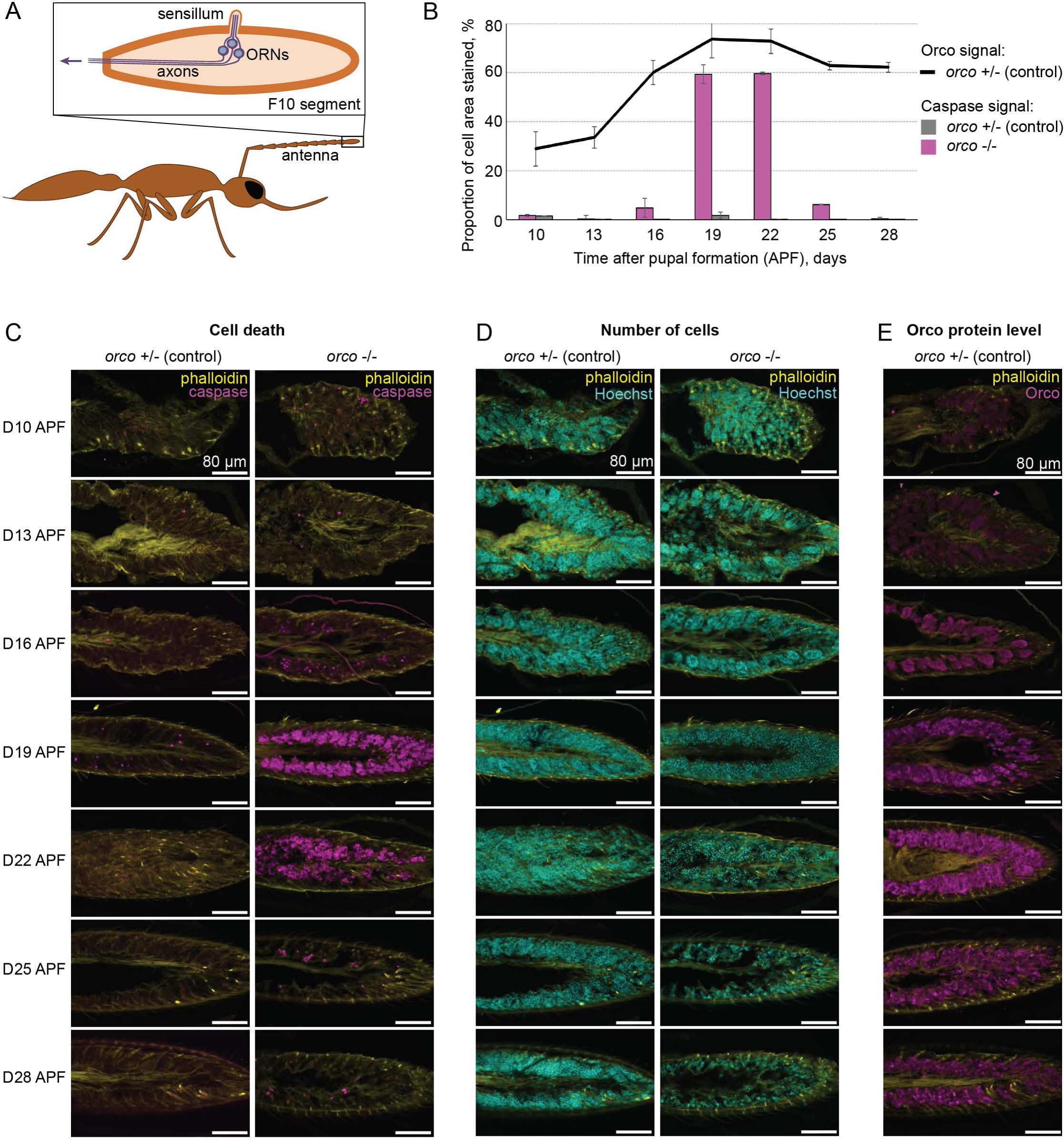
Temporal relationship between cell death and *orco* expression. **(A)** A simplified diagram illustrating the internal antennal structure of the olfactory system in ants. ORNs = olfactory receptor neurons. **(B)** Quantification (mean ± SEM) of immunohistochemistry staining shown in (C-E). Signal was quantified in Fiji (ImageJ) based on percentage of cell area stained. Hoechst was used to dictate total cell area. **(C-D)** Representative immunohistochemistry images of sectioned pupal antennae demonstrating the relationship between genotype (*orco* +/- (left) and *orco* -/- (right)), age (D10 APF (top) through D28 APF (bottom)), and cell death. Cell death is shown by the apoptotic marker cleaved caspase- 3 (magenta) in (C), and temporal changes in cell density are shown via Hoechst (cyan) in (D). **(E)** Representative immunohistochemistry images of sectioned pupal antennae demonstrating the relationship between age (indicated as in C-D) and prevalence of Orco (magenta).

The architecture of the olfactory system in insects (predominantly studied in *Drosophila*) largely follows the rule of ‘one receptor, one neuron’, similar to vertebrates (*3*). Each ORN selectively expresses a single odor-sensing, or tuning, receptor from one of the three major gene families – *odorant receptors* (*Or*), *ionotropic receptors* (*Ir*), or *gustatory receptors* (*Gr*) (*4, 5*). While all vertebrate and nematode olfactory receptors are G-protein-coupled receptors (*6, 7*), tuning ORs in insects bind to an obligate co-receptor (Orco), forming ligand-gated ion channels (*8-12*). Most insect species, including *Drosophila*, have fewer than 100 *Or* genes (*4, 13*); however, this number is dramatically expanded in hymenopteran insects, a feature considered pre-adaptive to social evolution (*5*). Further expansion in ants (300–500 *Or* genes) may have facilitated the recognition of complex social cues. In *Drosophila*, all ORNs that express the same tuning *Or* project axons to the same glomerulus in the antennal lobe (AL) of the brain, where they synapse onto projection neurons that transmit information into the central brain (*3, 5*). Consistent with the expansion of *Or* genes in ants, the number of AL glomeruli has also increased from 60 in *Drosophila* to 270–500 in female ants (*14-16*). In summary, ants have an expanded number of both *Or* genes and AL glomeruli, but it is unclear whether the same developmental paradigms as in *Drosophila* exist in ants to control the production of this expanded array of ORNs.

Until recently, studies of the biological function of ORs and ORNs in eusocial insects were limited by a lack of genetic tools. This has been overcome by the development of *orco* mutant ants via CRISPR/Cas9 and subsequent phenotypic analyses (*15, 16*). We have previously found that *orco* mutation in the jumping ant *Harpegnathos saltator* abolishes all OR-mediated olfactory sensation. Furthermore, mutant animals display communication deficits and abnormal behaviors, such as ‘wandering’ outside the nest, antennal twitching in the absence of an external stimulus, and inability to communicate with conspecifics (*16*). Surprisingly, unlike in *Drosophila* where Orco is only required for the prolonged survival of ORNs in adults but not for their development (*17, 18*), the loss of Orco in *H. saltator* dramatically reduces the number of ORNs in antennae and the number of AL glomeruli in newborn ants (*16*). The same phenotypes have been found in the clonal raider ant *Ooceraea biroi* (*15*) and the honeybee *Apis mellifera* (*19*), showing that social hymenopterans with their expanded *Or* repertoire require Orco for proper ORN development.

The exact sequence of developmental events and the underlying molecular mechanisms leading to the reduced numbers of antennal cells and AL glomeruli observed in *orco* mutants remain unknown. Are these phenotypes a result of ORN apoptosis or other developmental events? If the former, is it due to the absence of neuronal activity or to improper receptor trafficking (*20, 21*)? We show that all *Or*-expressing ORNs undergo massive apoptosis when Orco is absent. Only a small proportion of *orco*-expressing cells survive. Interestingly, these ORNs co-express genes encoding IR along with IR co-receptors (Ircos), suggesting that the activity of a functional receptor-coreceptor complex may be required for ORNs to survive during development. In addition, we found that some *Or* genes are also expressed in non-neuronal antennal cells that do not need Orco to survive. This *Or* gene expression might be spurious due to the leaky regulation of a neighboring gene in these cells.

## RESULTS

### ORNs undergo apoptosis during pupal development in *orco* mutants

To address the events leading to the reduction in antennal ORNs in adult *orco* mutant ants, we performed immunostaining of antennae across pupal development. The duration of pupation in *H. saltator* is approximately 30 days (*22*). Homozygous *orco* mutant and heterozygous control (*16*) pupae were harvested at 7 developmental stages: D10 D13, D16, D19, D22, D25, and D28, representing 10-28 days after puparium formation (APF). Their antennae were immunostained with antibodies against cleaved caspase-3 as a marker of programmed cell death (apoptosis), in conjunction with the nuclear marker Hoechst to confirm the loss of ORNs in late-stage pupae. In *Drosophila*, apoptosis of some ORNs normally occurs during neurogenesis (22-24 hours APF, which roughly translates to D7 APF in *H. saltator*) in order to eliminate extraneous ORNs during asymmetrical divisions of sensory organ precursor (SOP) cells (*5, 23-26*). Consistently, in heterozygous ant pupae, we observed occasional apoptosis (∼0-2% of total cells in the antenna) at the earliest time point examined (D10 APF), and a second small increase in the number of apoptotic cells at D19 (Fig. 1B, C), although the cause remains unclear. In the early days of development, homozygous pupae resembled heterozygotes closely in patterns of cell death, only beginning to diverge around D16. In *orco* mutant pupae, apoptosis greatly increased and was much more prevalent than their heterozygous counterparts, peaking between D19 and D22, and then ceasing around D25-D28 (Fig. 1B, C, D). Within the antennae, neuronal cells can be differentiated from other cell types based on location (deep below the cuticle, around the central axon bundle) and the shape of their nuclei (round and in cell clusters). Furthermore, neuronal cells (especially ORNs) make up the majority of antennal cells. Based on these features, we could infer that the dying cells are ORNs. In summary, a massive wave of apoptosis in mid-pupation (D19 and D22) leads to a greatly reduced ORN population by the end of pupation.

We also tracked the temporal pattern of Orco protein expression in developing heterozygous pupae. In pupae younger than D16, Orco stained faintly (Fig. 1B, E). Then the intensity of Orco staining greatly increased and peaked at D19 (Fig. 1B, E). Orco stained cells in grape-like clusters from D10 through D16 (Fig. 1E). Beyond this period, the cell clusters became larger while the boundary between cells was less obvious (Fig. 1D, E). Therefore, the massive apoptosis in homozygous mutant pupae (D19-D22) occurs after a dramatic increase in Orco protein in the control pupae.

### ORNs express *Or* genes before cell death in *orco* mutants

After establishing the correlation between ORN apoptosis and the increase of Orco, we asked two questions aiming to further understand the developmental context of these events: (1) Does the cell death promptly follow neurogenesis or does it occur at a later developmental stage? (2) Are tuning *Ors*, like *orco*, expressed before the initiation of cell death? To address these questions, we performed bulk RNA-sequencing (RNA-seq) on WT and mutant antennae at D10, D15, D20, and D25 APF.

Principal component analysis (PCA) showed that developmental stage was the primary source of variance in gene expression (Fig. 2A). We identified that “early genes” (expression peaking at D10) were enriched for mitotic terms, “intermediate genes” (expression peaking at D15) were enriched for terms associated with differentiation of neurons and non-neurons, as well as neuronal activity, and “late genes” (expression plateauing at D20) were enriched for metabolic terms (Fig. 2B-C, Fig. S1A-C). These data suggested that the peak of cell death in *orco* mutants observed at D19 and D22 occurred long after SOP division (unlike the programmed cell death in dividing ORNs in *Drosophila*).

**Figure 2.**
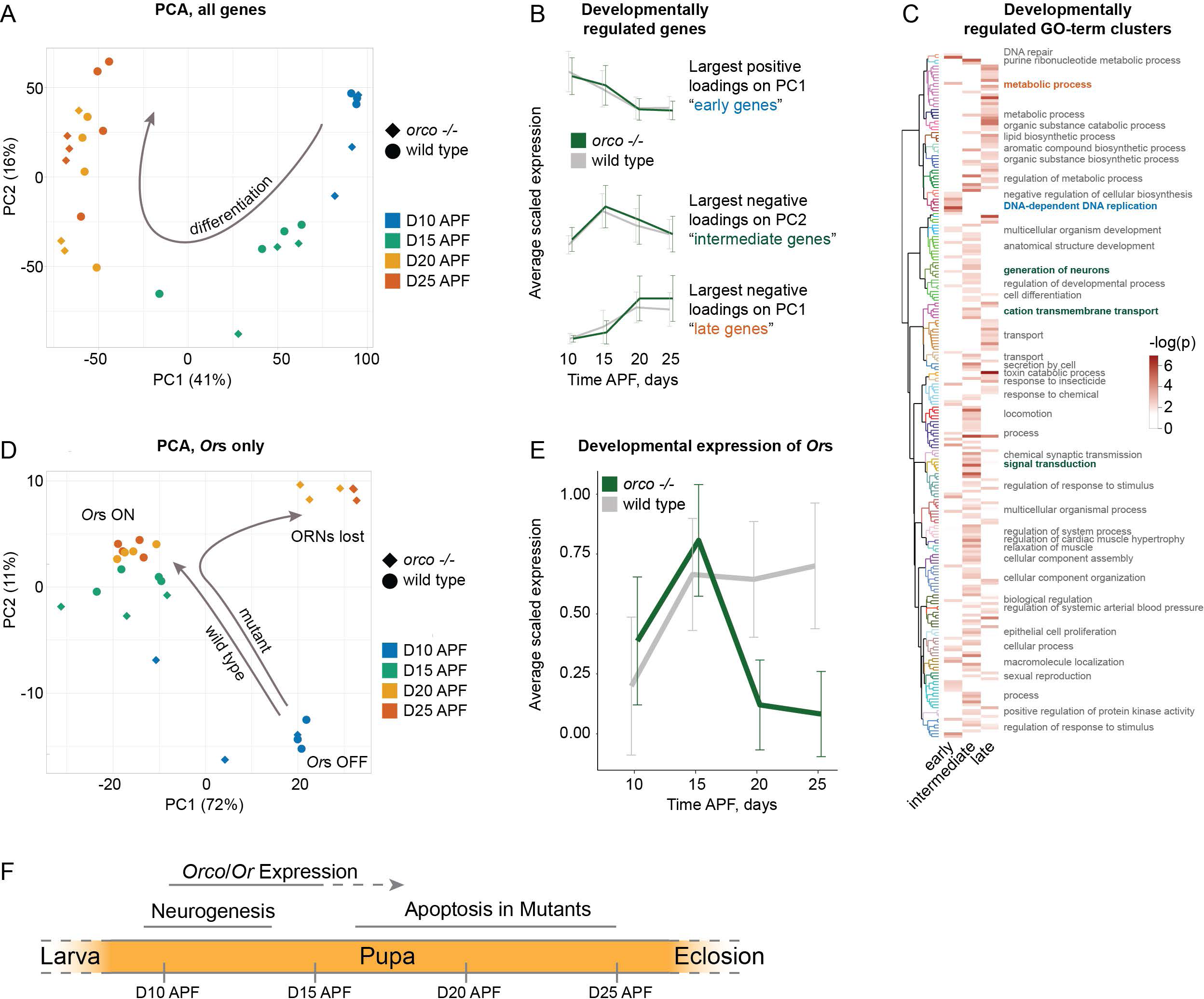
Developmental trajectory in wild type and *orco* mutant antennae. **(A)** Whole transcriptome-based PCA of bulk RNA-seq samples collected from wild-type and mutant pupae of different ages. Percentages represent the amount of variance explained by each PC. The arrow illustrates the developmental trajectory from D10 to D25. **(B)** The identification strategy and the expression pattern of the inferred “early”, “intermediate”, and “late” gene sets. The line represents average expression across genes after scaling the expression of each gene to its maximum expression value. The error bars represent standard deviation in the scaled expression values across genes. **(C)** The heatmap shows minus logarithm of the enrichment p-values for every GO-term significantly enriched in a minimum of one gene set. The dendrogram on the left represents grouping of the GO-terms by their semantic similarity, and branch color corresponds to cluster identity. The labels on the right are cluster identities. **(D)** PCA based on *Or* genes only. The arrows illustrate wild-type and mutant developmental trajectories. **(E)** The expression pattern of *Or* genes. The line represents average expression across genes after scaling the expression of each gene to its maximum expression value. The error bars represent standard deviation in the scaled expression values across genes. **(F)** A summary of olfactory developmental process in *H. saltator*: SOP division is ongoing during D10-D13 (APF); *Or* and *orco* mRNAs are visible at D10 and reach their peaks at D15, and Orco protein level increases from D10 and peaks at D19; in the mutant, apoptosis starts at D16, peaks at D19-D22 (thicker line) and fades at D25, later than SOP division and *Or/orco* expression.

For *Or* genes, both PCA (Fig. 2D) and developmental expression profile of *Or* genes (Fig. 2E) showed that expression of *Ors* started at D10, increased dramatically between D10 to D15 and remained largely stable after D15 in WT pupae. In *orco* mutant pupae, however, although a similar pattern was observed at D10 and D15, *Or* expression dropped sharply between D15 and D20 and remained low at D25 (Fig. 2E). A parallel pattern was observed with *orco*, which comprised a deletion of two nucleotides (*16*): its mRNA level increased from D10 to D15, and then became stable in the WT, while it dropped sharply after D15 in the mutants (Fig. S1D). These data allowed us to infer the following developmental sequence: cell proliferation occurs around D10, followed by ORN differentiation at D15-D16 with peak expression of *orco* and *Or*s. Concurrently, apoptosis commences in the *orco* mutants, and then intensifies dramatically, peaking at D19-D22 (Fig. 2F). The death of cells leads to a sharp decline in the expression of tuning *Or*s in the mutants. In summary, massive apoptosis occurs in differentiating neurons that express *Ors* and *orco*.

### Co-expression of functional *Ir* complex genes rescues *orco*-expressing ORNs

To gain insight into what cell types undergo apoptosis in *orco* mutants, we performed single-nucleus RNA-sequencing (snRNA-seq) of WT and mutant antennae. Our data set contained ∼16k nuclei, including ∼13k WT nuclei and ∼3.5k nuclei from the mutant (Fig. 3A, B). We first observed that neurons made up strikingly different proportions of the total number of nuclei in WT (67%) and mutant samples (12%). Such dramatic underrepresentation of neurons in mutants indicated that neurons are disproportionately affected by cell death, consistent with our caspase staining. Thus, we set out to investigate the specific types of neurons that were absent in the mutants (Fig. 3C). We classified individual neuronal cells based on the repertoire of receptors they expressed, specifically *Gr*s, tuning *Ir*s, *Ir* co-receptors (*irco*s), tuning *Or*s, *orco*, an ammonia transporter *Rh50*, and a mechanoreceptor *nompC* (see methods section for details). This allowed us to place the neurons into categories as shown in Fig. 3C, e.g. “mechanosensory neurons” or “chemosensory neurons expressing *Gr*(s) or *Ir* and their co-receptor(s)”. Notably, we observed several cell types co-expressing *irco*s with *orco* and tuning *Or*s. In *Drosophila*, an *irco* gene *Ir25a* is expressed in most *Or/Orco*-positive ORNs. In contrast, the two ant orthologs of *irco Ir25a* (*HsIr25a.1* and *HsIr25a.2*) were only expressed in *Ir*, *Gr,* and *nompC*-positive neurons. In contrast, another *irco* – *HsIr8a* – was co-expressed with some *Orco/Or*-positive ORNs (Fig. 3D). The fourth *irco* gene, *HsIr76b*, displayed a similar expression pattern to that of *HsIr25a.1* and *HsIr25a.2* with the exception of a peculiar cluster of *orco*-positive cells that co-expressed it with *orco, HsIr8a,* and a tuning *Ir* (see below).

**Figure 3.**
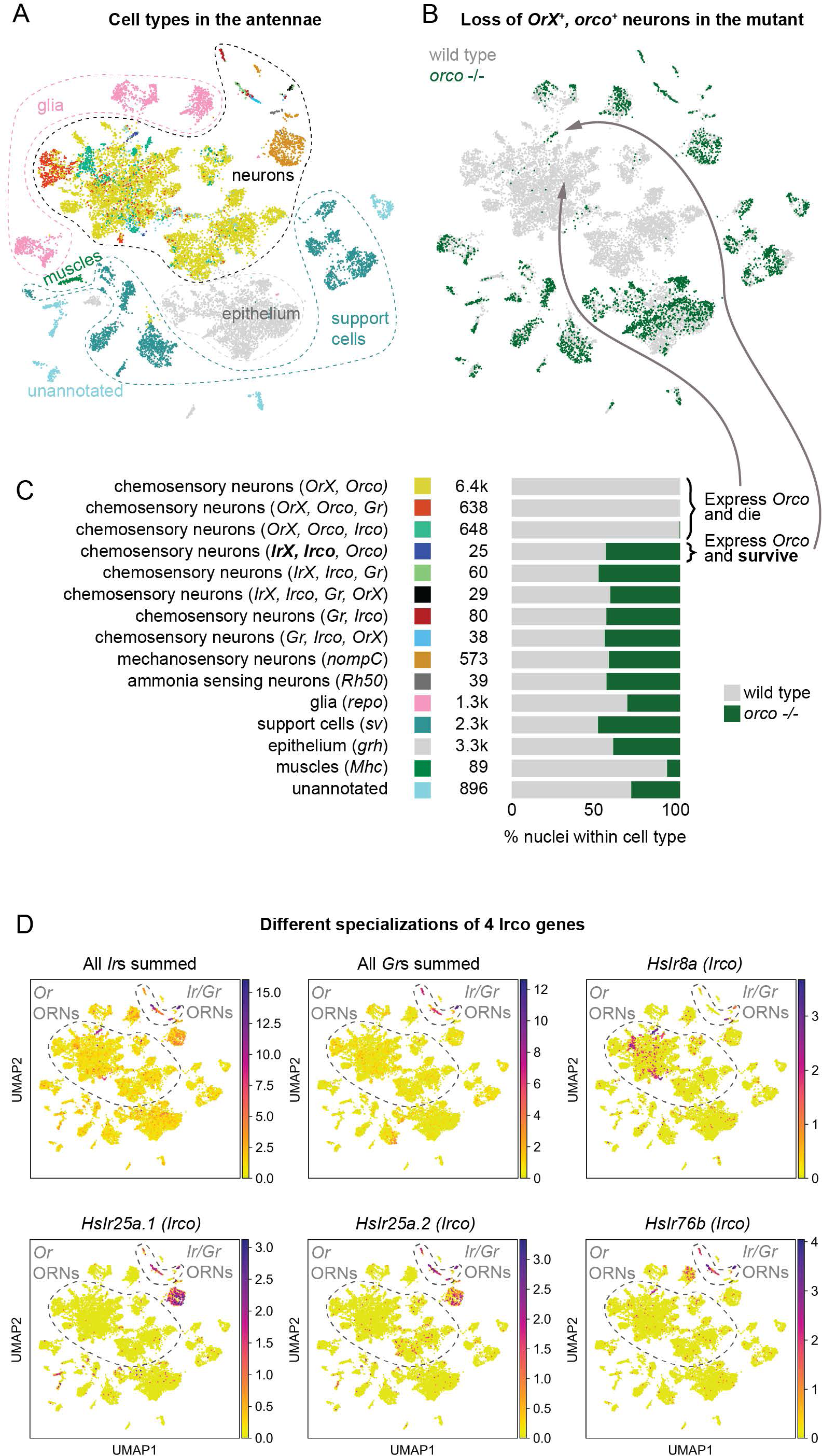
Cell death at single-cell resolution. **(A)** Combined UMAP plot showing the distribution of different cell type categories from (C). **(B)** The same UMAP plot as in A, but with color showing WT and mutant cells. **(C)** The list of cell type categories used in this study. Category labels are based on expression data only and should not be interpreted as functional descriptions because some neurons may be polymodal, and molecules traditionally considered chemoreceptors may have functions beyond the reception of chemical signals. Colored rectangle near each category shows which color was used for nuclei of this category in A. The number is the total number of nuclei from the category in the combined data set. The stacked bar plot shows the proportion of WT and mutant nuclei from the category in the combined data set. **(D)** Combined expression of all *Ir*s, all *Gr*s, and the expression of the four *irco* genes plotted on the UMAP.

There were almost no mutant nuclei belonging to the *Or/orco*-expressing ORNs (*OrX*, *orco*) (Fig. 3B, C), indicating that Orco is necessary for the survival of the ORNs that express tuning *Ors*. Neurons that expressed *Gr*(s) or *irco*(s) in addition to a tuning *Or* and *orco* were, likewise, almost completely absent from the mutant sample. Thus, the additional expression of these receptors or co-receptors was not sufficient to rescue the effect of the *orco* mutation. However, neurons that expressed both tuning *Ir*(s) and *irco*(s) in addition to *orco* alone fully survived in the mutants (Fig. 3B, C). This showed that Orco is not necessary for the survival of ORNs that express tuning *Ir*s and *Irco*s. Finally, neurons that did not express *orco* and non-neuronal cells were not affected in the mutants, with a surprising but currently unexplained exception of muscle cells, which were reduced in the absence of Orco (Fig. 3C). In summary, we observed that the antennae of *orco* mutants specifically lacked ORNs expressing tuning *Or*s and *orco*, including those cells that additionally expressed *Gr*s or *irco*s without tuning *Ir*s. In contrast, some ORNs expressing *orco* were maintained in the mutants as long as they additionally produced a functional IR/Irco complex.

### Non-neuronal cells expressing *Or*s without *orco* remained in *orco* mutants

We also performed bulk RNA-seq on the antennae of adult WT, heterozygous, and homozygous *orco* mutants. Having established that most *Or*-expressing ORNs die, we expected to see a drastic reduction in *Or* expression in the mutant antennae. This was indeed the case for most *Or*s in the homozygous mutants (Fig. 4A). Unexpectedly, several *Ors* not closely related to each other phylogenetically, such as *HsOr370, HsOr61*, *HsOr247*, and *HsOr202*, retained considerable expression in the mutants (Fig. 4A). We sought to confirm this observation using our snRNA-seq data (Fig. S3A). Surprisingly, plotting the expression of these genes on the UMAP revealed that these *Or* genes were expressed both in neurons and in various populations of non-neuronal cells including specific subtypes of glia and support cells (Fig. 4B, C, Fig. S3C). Thus, only their ORNs died while non-ORN cells survived in *orco* mutants, which explained why they remained detectable. In mammals, some *Or* genes exhibit expression outside olfactory cells, and *Or* genes with such expression patterns are located near non-*Or* genes expressed in the same non-olfactory tissues (*27-29*). Accordingly, we found that non-*Or* genes neighboring *HsOr370, HsOr61*, *HsOr247*, and *HsOr202* exhibited expression in the same non-neuronal tissues as these *Or*s (Fig. 4C). In some clustered *Or*s, we noted a pronounced inverse relationship between the strength of the non-neuronal expression and the physical distance to the non-*Or* gene: *HsOr202*, which is the closest to the glia-expressed non-*Or* gene *LOC105190781*, had the strongest expression in glia, while more distant *HsOr201* and *HsOr200* had weaker expression in glia (Fig. 4C, Fig. S3C). Additionally, *Or* expression in non-neuronal tissues tended to be lower than in ORNs (Fig. S3B), consistent with the idea that this “ectopic” expression is driven by the regulatory elements of neighboring genes. We proceeded to systematically analyze *Or* expression in the mutant and found many additional cases of expression in non-neuronal tissues as well as several cases where *Or* genes were expressed in mechanosensory neurons, which also survived in the mutant (Table 1). Of note, none of these non-ORN cell types expressed *orco*. In summary, the *orco* mutants lost the expression of the majority of *Or* genes, while retaining the expression of some *Ors* in certain non-neuronal cells (such as support cells and glia) or non-olfactory neuronal cells (mechanosensory neurons). This expression is independent of *orco* and may be driven by closely located regulatory elements of non-*Or* genes, leading to leaky expression of the *Or* genes (*30, 31*). A majority (∼70%) of *Or* expression in the non-ORNs can be explained by the leaking regulation of neighboring genes, while ∼30% of *Or* expression cannot (Table 1) (see Discussion).

**Figure 4.**
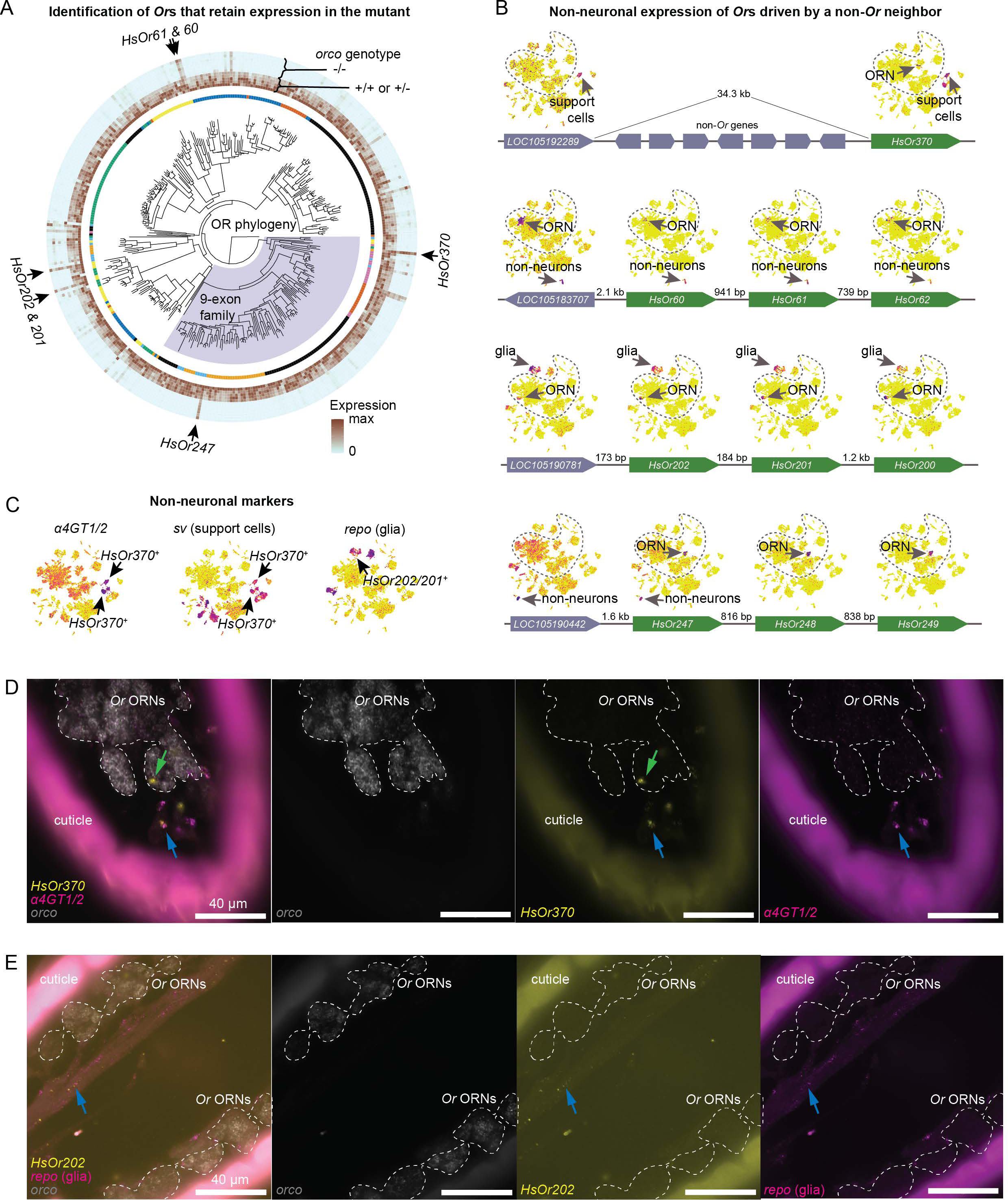
Identification of Orco-independent ORs and their non-neuronal expression. **(A)** In the middle, the phylogeny of *Or* genes with the pheromone-sensing 9-exon *Or*s shaded blue. The inner circle depicts the genomic location of each gene, such that all genes residing on the same scaffold are colored in the same way. The outer circles show scaled expression in WT/heterozygote and mutant samples. Arrows point at genes whose expression is most prominently maintained in the homozygous mutant. **(B)** UMAP plots showing the expression of the Orco-independent *Or*s that are marked with arrows in panel A and their neighbors. Dashed line encircles neurons. The genomic structures of the corresponding loci are depicted under the UMAP plots. Green boxes correspond to *Or* genes and grey boxes correspond to non*-Or* genes. The arrow at the end of each box points to the 3’ end. Distances are between the closest ends of two gene predictions in the HSAL60 annotation set. **(C)** UMAP plots showing the expression of selected tissue markers. **(D)** Representative HCR RNA-FISH images of sectioned WT adult antennae. Here, we show the relationship between an Orco-independent gene (*OR370*, yellow), identified support cell marker *α4GT1/2* (magenta) and a neuronal marker (*orco*, grey). Green arrows show cases of *OR370* expression in neuronal cells. Blue arrows demonstrate cases where *OR370* co-localizes with support cell markers, but not the neuronal marker. **(E)** Representative HCR RNA-FISH images of sectioned WT adult antennae showing the relationship between an Orco-independent gene (*Or202,* yellow) and glial cells (*Repo*, magenta). The blue arrow indicates a case of co-localization of the two markers within the antennal axon bundle, independent of our ORN marker *orco* (grey).

**Table 1.**
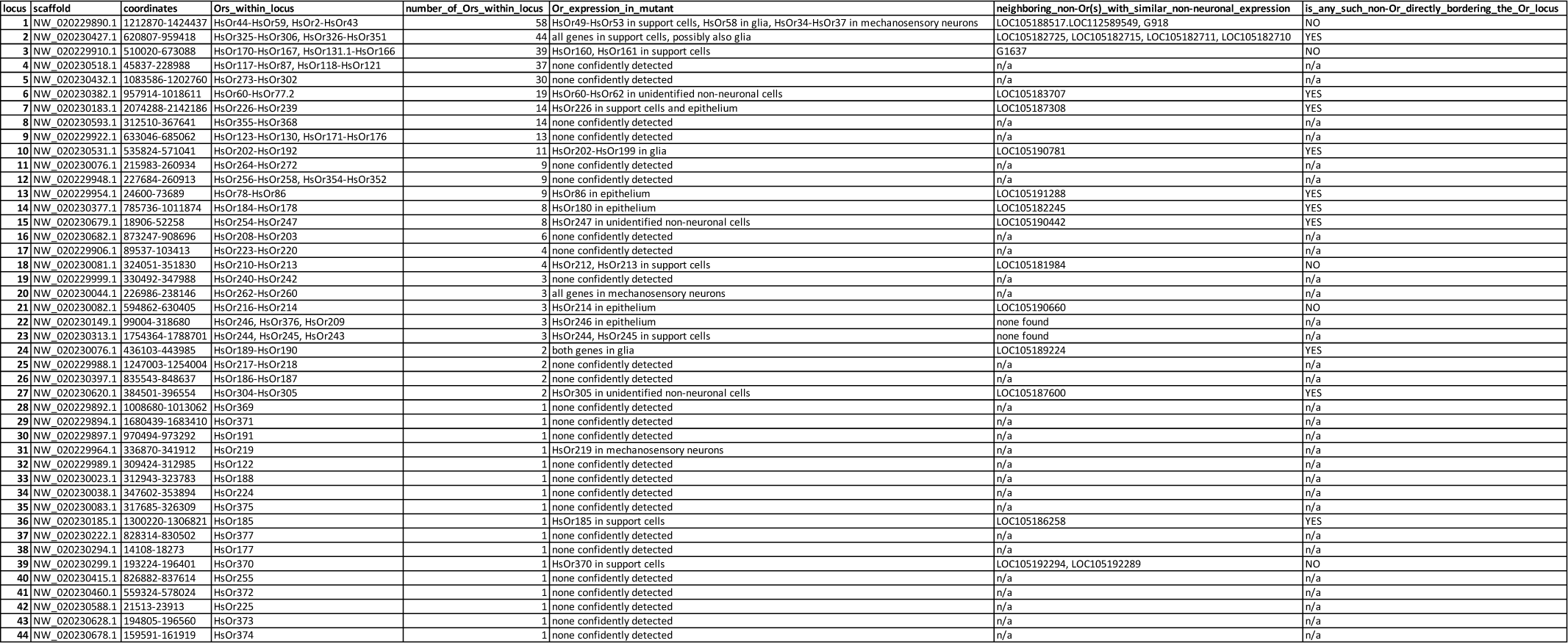
List of *Or* loci, the description of their expression pattern in mutant snRNA-seq data, and neighboring non-*Or* genes which could potentially drive non-neuronal expression of *Or*s.

We thus checked whether other insects (e.g. *Drosophila*) also exhibit expression of chemoreceptor genes in non-neuronal antennal cells. We accessed the antennal snRNA-seq data generated as part of the Fly Cell Atlas initiative (*32*) and visualized the expression pattern of all *Gr*, *Ir*, and *Or* genes on the UMAP. In contrast to *H. saltator*, no *Or* was expressed outside ORNs in *D. melanogaster*. However, we identified four *Ir* genes with non-neuronal expression. Specifically, *Ir41a* was expressed in *sv*-positive populations likely representing support cells, and *Ir47a, Ir47b,* and *Ir60a* were expressed in different populations of glia as identified by the expression of *repo* (Fig. S4). Similar to the Orco-independent expression of the non-neuronal *Or*s in *H. saltator*, the non-neuronal cells expressing *Ir* genes in *D. melanogaster* did not express any *irco*. Thus, *D. melanogaster* also exhibits co-receptor-independent expression of chemoreceptors in support and glial cells of the antennae, although the chemoreceptors expressed are not *Or*s as in *H. saltator*.

To explore the localization of *Or*s in adult antennae, HCR RNA-FISH probes were designed for *HsOr206* (an *Or* gene solely expressed in ORNs) and *HsOr370* (an *Or* that is expressed in non-neuronal cells in addition to ORNs). The Orco-dependent *Or* was always expressed with neuronal markers (i.e., *Syt1* and *orco*); however, the Orco-independent *Or* was also expressed in apparently non-neuronal cells, which exhibited elongated nuclei and were primarily found adjacent to the cuticle (Fig. S3D). Our snRNA-seq data showed that *HsOr370* was expressed in two clusters of *sv*-expressing support cells (Fig. 4B, C). We selected marker genes and designed probes for these clusters. Two of the selected genes, *α4GT1/2 (LOC105190591)* and *LOC105182650*, were expressed in both *HsOr370*-positive clusters, while *LOC105188530* was only expressed in one of them (Fig. 4C, Fig. S3E). Cells expressing these genes generally appeared just under the cuticle. *HsOr370* was also expressed in *orco*-expressing ORNs (Fig. 4B, Fig. S3F), thus exhibiting expression in both neuronal and non-neuronal cells. Unlike our other support cell markers, *LOC105182650* stained cytoplasm. The unusual tubular shapes of these cells further confirmed their cell type (Fig. S3G). Furthermore, the cytoplasm of these cells seemed to project through the pores of the cuticle towards the sensilla. The localization and shape of our identified non-neuronal cells suggested that they could play a role in supporting the function of ORNs (*33*). In addition, HCR RNA-FISH allowed us to confirm the expression of *HsOr202* in glial cells (Fig. 4C) that were present around antennal ORN axons (Fig. 4E, blue arrow) via co-localization with the glial marker *repo*.

## DISCUSSION

We and others previously found that null mutations in *orco* lead to a wide range of neuronal, physiological, and behavioral defects in ants and honeybees (*15, 16, 19*). In this study, we further reveal that developing ORNs undergo massive apoptosis in *orco* mutant ants, providing a cellular mechanism underlying their defective neural development.

ORNs start to die at D16 APF in mutant ants, with apoptosis peaking at D19 and D22. The massive apoptosis occurs after the onset of expression of *orco* and *Or* genes (*orco* mRNA peaks at D16, and Orco protein plateaus at D19). Thus, there appears to be a developmental switch around D15-19: at this point, Orco abundance increases to reach its peak, and it simultaneously becomes essential for the survival of ORNs (Fig. 2F). As a note, *orco* mutants exhibit a drastically reduced number of glomeruli in the antennal lobe (*16*): only ∼20% of glomeruli remain. This can be explained by the lack of axon projection due to ORN cell death, suggesting that axons from the majority of ORNs have not yet projected to glomeruli when massive ORN apoptosis occurs in the mutants (D19-D22) (Fig. 2F). This suggests that *Or* genes are expressed before axon targeting in *H. saltator*, consistent with the evidence in another ant *O. biroi* (*34*).

Why might Orco be required for the survival of developing ORNs? Orco itself and Orco/OR complexes may generate spontaneous activity during ORN development, as in *Drosophila* (*35-38*), and activity-dependent survival of neurons is essential for refinement of neuronal network during neural development and for removal of extra cells produced in neurogenesis that are not functional (*39*). Loss of activity is known to affect the survival of developing ORNs in mice (*20, 40*). In ants, loss of Orco may lead to loss of spontaneous activity, and thus induce ORN apoptosis. A small proportion of *orco*-expressing ORNs that co-express *Ir* and *irco* survive in the mutant ants, suggesting that IR/Irco complexes might also generate spontaneous activity which compensates for loss of Orco-induced activity. If spontaneous neuronal activity is required for the survival of ORNs in ants, why is this different in *Drosophila* (*17*)? In *Drosophila*, SOPs undergo two rounds of cell division, generating up to four ORNs in each sensillum. During this neurogenesis, a proportion of extra neurons differentially specified by Notch are eliminated (*23, 26*). Although this early apoptosis has not been analyzed in ants, it is tempting to speculate that a low level of apoptosis found at early pupal stage (D10) is driven by Notch-dependent cell specification. It is important to note that each ant sensillum contains up to 130 ORNs (*41*), suggesting that SOPs divide many more times than in *Drosophila*, which might generate many cells. Therefore, spontaneous activity might be utilized in ants as an additional mechanism to remove extra cells produced during neurogenesis. Interestingly, there is an increase in apoptotic cells at D19 in WT pupae, coinciding with the occurrence of massive cell death in the mutants. Whether this is driven by lack of neuronal activity merits further investigation.

In *Drosophila*, Orco is required for the trafficking of OR proteins to dendrites (*38*). Disrupted trafficking might lead to an excess of OR protein in the endoplasmic reticulum (ER) of ant ORNs, which in turn causes ER stress and may induce apoptosis. Two lines of evidence including (1) the death of all tuning *Or*-expressing ORNs and (2) the survival of ORNs that express *orco,* tuning *Ir* and *irco*, but no tuning *Or*, are consistent with this hypothesis as ER stress would be triggered in the former but not the latter. However, some *Or*s are expressed in non-neuronal cells that do not express *orco*. This seems inconsistent with apoptosis induction by ER stress, although *Or* expression in non-neurons tends to be lower than their neuronal expression, and the non-neuronal cells might exhibit other properties rendering resistance to ER stress.

We have also characterized a mechanism for the survival of *Or*-expressing cells in mutant ants. For instance, *HsOr370* is expressed in both neuronal and non-neuronal cells. The latter are likely support cells located near the antennal cuticle which surround the dendrites of ORNs, and these cells do not die in the *orco* mutant. There are 3 types of support cells as identified in *Drosophila*: thecogen (sheath) cells, trichogen (shaft) cells, and tormogen (socket) cells (*42, 43*). It remains unclear which cell type expresses *HsOr370*. *HsOr202* is co-expressed with *repo*, a glial cell marker. Interestingly, when we reanalyzed the published data from *Drosophila* snRNA-seq (*32*), we found that a few *Ir* but no *Or* genes are expressed in glial and support cells. As these cells are derived from the same developmental lineage as ORNs (*32, 42*), it is tempting to speculate that certain *Drosophila Ir* and *Harpegnathos Or* genes are turned on in non-neuronal cells under the same regulatory process as in ORNs. Leaky *Or* expression has been observed in mammals, where *Or* genes are co-expressed with neighboring non-*Or* genes in non-neuronal cells (*27*). Indeed, we found (1) similar spatial expression profiles between *Or* genes and their non-*Or* neighbors; (2) gradually faded expression of *Or* genes in the *Or* cluster depending on their distance to the non-*Or* gene expressed in these cells, consistent with the notion of leaking expression. However, some *Or* expression cannot be explained by their direct neighboring genes. It is possible that these *Or* genes are regulated by distant enhancers due to high-order chromatin structure.

Mounting evidence also revealed the role of chemosensory receptors expressed in non-neuronal cells. In mammals, non-neuronal ORs play a role in development, reproduction, and immune response (*44-46*). In *Drosophila*, certain gustatory receptors – such as *Gr28* – and ionotropic receptors – such as *Ir25a* (*irco*) and *Ir21a* (tuning *Ir*) – are expressed in non-chemosensory neurons and mediate temperature sensing (*47-49*). In mosquitoes, *orco* and some *Or*s are expressed in testes and sperm cells and may be involved in sperm chemotaxis (*50*). However, the role of ant *Or*s and *Drosophila Ir*s in glia and support cells remains unclear. As ant *Or* genes expressed in non-ORN cells do not co-express *orco*, it is possible that these ORs are not functional or their functions are independent of olfaction.

Co-expression of chemosensory receptors is a common phenomenon in *C. elegan*s, which has many more *Or* genes than sensory neurons (*51*); however, *Or* expression in insects and vertebrates largely follows the rule of ‘one neuron, one receptor’ (*3, 5*). Recently, exceptions have been found in mosquitoes and *Drosophila,* where some neurons express multiple *Or* genes. *orco* is also expressed in neurons expressing *irco*s (*52-54*). Consistently, we found a wide range of co-expression between different classes of chemosensory receptors in ants. In contrast to dipteran insects where *Ir25a* is commonly co-expressed with *orco*, the main co-expression pair in ants is *orco*-*Ir8a*. Although the precise function remains unclear, the co-expression might provide evolutionary advantages: for example, an IR complex could modulate neuronal activity by altering membrane resistance (*54*) or allow a limited number of neurons to detect many chemical cues from the same target (*52*).

In summary, our study revealed the temporal windows of SOP cell division, as well as patterns of *orco/Or* gene expression and neuronal survival during ant ORN development, consistent with spontaneous activity generated by functional receptor complexes in the developing neurons. Activity-dependent survival may represent a cellular mechanism in ants to generate the correct number of ORNs and to form a proper olfactory map. Hymenoptera have expanded their *Or* gene repertoire, which possibly resulted in unique developmental events among insect chemosensory systems to support eusociality.

## MATERIALS AND METHODS

### Experimental design and model

Animals for HCR and immunostaining were maintained and collected at the UF Department of Biology. Heterozygous females were obtained by crossing mutant males with wild-type females. Heterozygous and homozygous mutant females used in this study were collected from the offspring of heterozygous females paired with mutant males. Newly pupated individuals were painted with a single colored dot near the posterior end of the abdomen to mark the day on which they underwent pupation. Pupae were then assigned an age at which to be harvested (10-28 days after puparium formation). Prior to harvesting, sex was confirmed by examining the body shape of the pupa. Females could be identified by their long mandibles, large heads, and stout bodies. Males, identified by their long antennae, small heads, and narrow bodies, were removed from the study. To prepare pupal antennae for experiments, the pupal casing was removed by clipping a small hole near the base of the pupa’s abdomen, and gently pulling the pupa out of its casing from the posterior end. This prevented any potential damage to the antennae, which were then clipped at the base. Bodies were stored at -80°C and later used for genotyping by Sanger sequencing of the mutation site. Antennae were either used immediately or stored in O.C.T. at -80°C after fixation in paraformaldehyde. To prepare adult antennae for experiments, callow workers (1-2 days post-eclosion) were collected from wild-type colonies. Antennae were clipped at the base and were immediately used for experiments.

Wild-type animals used for snRNA-seq were collected at NYU Grossman School of Medicine. Six (library 1) or 26 (libraries 2 and 3) adult females of unknown age were collected from stock colonies. Their antennae were clipped at the base and processed for nuclei extraction immediately. *Orco* mutant animals for snRNA-seq were collected at the University of Florida. Homozygous females reach adulthood, but they generally die early and do not leave any progeny, so the mutation is propagated in the lab by selecting for heterozygous reproductive females. To collect homozygous mutants for the experiment, antennae of the freshly eclosed female progeny of heterozygous females were clipped at the base, and both the antennae and the remaining bodies were placed at -80°C for storage. These samples were shipped to NYU Grossman School of Medicine on dry ice. DNA was extracted from the remaining bodies and genotyped by Sanger sequencing of the mutation site. Afterwards, the antennae of seven individuals that were found to be homozygous mutants were removed from storage and processed for nuclei extraction.

### Immunostaining of pupal antennae

The protocol for immunostaining, adapted from a previous protocol (*16*), is described here. Pupal antennae were fixed in 4% PFA diluted in 1X PBS with 0.3% Triton X-100 for 30 minutes, washed twice with 0.3% PBST, and underwent overnight incubation in 30% sucrose at 4°C. Sections were taken the following day at a 15um thickness, with focus on the most distal portion of the antennae (F5-F10). If whole antennae are difficult to section, the scape and F1-F4 may be removed prior to sectioning. Tissue was fixed on slides using 4% PFA (as prepared previously) for 30 minutes at room temperature before two washes with 0.3% PBST. The tissues were then incubated overnight at 4°C in a primary antibody solution (1:400 primary antibody from rabbit). After ensuring all primary antibody is removed from the slide via two 0.3% PBST washes, tissues were incubated at room temperature for 2 hours in a secondary antibody solution containing Alexa Fluor 555 anti-Rabbit secondary antibody (1:400, Thermo), Alexa Fluor 488 Phalloidin (1:200, Thermo), and Hoechst (1:1000, Sigma) before being mounted for microscopy.

### HCR RNA-FISH of adult antennae

Our protocol for HCR RNA-FISH was adapted from several established previously (https://www.molecularinstruments.com/, (*52, 55*)) and is described here. Antennae were harvested from callow female workers (1-2 days post eclosion) and fixed in 4% PFA diluted in 1X PBS with 0.1% Triton X-100 for 30 minutes, washed twice with 0.1% PBST, and underwent overnight incubation in 30% sucrose at 4°C. Sections were taken at a 10um thickness and fixed on slides using 4% PFA (as prepared previously) for 30 minutes at room temperature. Tissue was then dehydrated and subsequently rehydrated with a graded series of 5-minute MeOH/0.1% PBST washes prior to incubating in a 10 µg/mL proteinase K solution for 10 minutes. Slides then underwent a 10-minute incubation in probe hybridization buffer warmed to 37°C. Chambers were made for each slide consisting of a coverslip with double-sided tape lining two parallel edges to raise the coverslip slightly after application to the slide. 1.6pmol of each probe set in 100ul warmed probe hybridization buffer was added beneath the raised coverslip chamber. The chamber was sealed using rubber cement, and the slides incubated overnight at 37°C. Probes were removed using a graded series of 15-minute washes with probe wash buffer and 5X SSC with 0.1% PBST before incubating at room temperature for 30 minutes in amplification buffer. For each amplifier set, 6pmol of hairpin h1 and 6pmol of hairpin h2 were prepared by heating at 95°C for 90 seconds and cooling to room temperature in a dark space for at least 30 minutes. These were quickly added to 100ul of room temperature amplification buffer. Again, coverslip chambers were created and added to slides, and the amplification buffer mixture was added before the chamber was sealed with rubber cement and the slides left overnight to incubate at room temperature in a dark space. The amplification buffer mixture was removed via two 30-minute washes with 5X SSC and 0.1% PBST before incubation with Hoechst (1:1000) for 15 minutes. Following two subsequent 30-minute washes with 0.1% PBST, slides were mounted for microscopy using Vectashield mounting medium.

### Microscopy and image generation

All tissue samples were imaged using an Olympus IX81-DSU Spinning Disk confocal microscope at the University of Florida Interdisciplinary Center for Biotechnology Research. Z-stacks were acquired with 1um between each focal plane for all samples. Images were generated and colorized using Fiji (ImageJ).

### Bulk RNA-seq

Antennae for bulk RNA-seq were frozen with liquid Nitrogen and ground into a fine dry powder using a pestle. All bulk antennal RNA was extracted via a standard protocol using TRIzol™ Reagent, followed by ethanol precipitation. DNase treatment was performed in-solution, followed by a second round of TRIzol™-chloroform and ethanol precipitation to remove the treatment. To generate libraries, we used the NEBNext® Ultra™ II RNA Library Prep Kit for Illumina® with an input of 110 ng RNA and 12 PCR cycles. Pooled libraries were sequenced in several rounds using either HiSeq2500 or NovaSeq6000 sequencing systems.

### Nuclei extraction

Nuclei extraction protocol, largely adapted from fly and mosquito protocols (*32, 52, 53*), is described below. Dounce homogenizer, pluriStrainers, and tubes were pre-wetted with the corresponding buffer before adding the sample to prevent the adhesion of nuclei and to minimize sample loss.

1. Prepare a solution containing 1 % bovine serum albumin and 1 mM RNaseOUT in 1X phosphate buffered saline, pH 7.4 (PBS-BSA).
2. Prepare homogenization buffer:

3. Chill a metal cup on dry ice and chill a pestle by immersing it into liquid nitrogen.
4. If starting with fresh tissue, flash-freeze antennae in liquid nitrogen.
5. Empty freshly frozen antennae or antennae previously stored at -80°C into the cup and grind them while keeping the cup on dry ice.
6. Place the cup on wet ice until thawed.
7. Add 1 mL of homogenization buffer and wash down remaining sample from the pestle and the walls of the cup.
8. Transfer the entire sample into a Dounce homogenizer and release nuclei by 20 strokes of loose, 20 strokes of tight, 20 strokes of tight pestle (avoid foaming with steady consistent motion). Briefly chill on wet ice between the stroking series.
9. Split the suspension into two halves. Do the following for each half: filter the suspension through a 40 um Flowmi strainer directly into a 20 um pluriStainer inserted into a 1.5 mL tube.
10. Centrifuge both tubes for 10 min at 500 g at 4°C.
11. Without disturbing the pellet (which is not always formed), discard the supernatant.
12. Resuspend each sample in 250 uL PBS-BSA by pipetting 20 times.
13. Filter each sample three times through a 40 um Flowmi strainer and finally into a 10 um pluriStainer inserted into a 1.5 mL tube.
14. Combine the two sample halves.

| Component | Stock concentration | Amount | Final concentration |
| --- | --- | --- | --- |
| Nuclease-free water | - | Add to 10 mL final volume | - |
| Sucrose | - | 0.856 g | 250 mM |
| Tris, pH 8.0 | 1 M | 100 uL | 10 mM |
| Potassium Chloride | 1 M | 250 uL | 25 mM |
| Magnesium Chloride | 1 M | 50 uL | 5 mM |
| Triton X-100 | 100 % | 10 uL | 0.1 % |
| RNaseOUT | 100 mM | 50 uL | 0.5 mM |
| Protease inhibitor | 50X | 200 uL | 1X |
| DTT | 100 mM | 10 uL | 0.1 mM |

### Nuclei sorting and snRNA-seq library prep

Nuclear suspensions were stained with 5 ug/mL Hoechst and processed on the FACSAria II (BD Biosciences) cell sorter. First, single particles were gated in FSC-A vs FSC-W coordinates, and the resulting population was plotted in Hoechst vs FSC-A coordinates (Fig. S5A). Two or three subpopulations with varying and largely overlapping FSC-A signal but highly uniform and distinct levels of Hoechst fluorescence appeared on the plot, consistent with previous reports of nuclei with different ploidy in fly nuclear suspensions (*32*). All such subpopulations were included into the gate used for sorting. Additionally, a fraction of particles had variable but intermediate (higher than unstained control but lower than the nuclear “bands”) levels of Hoechst fluorescence. These particles were considered debris and were not included. Sorted nuclei were collected into a 1.5 mL tube containing 20 uL PBS-BSA. 43.2 uL (or less, if the volume was insufficient) of the final suspension was used as an input into scRNA-seq library prep using Chromium Next GEM Single Cell 3’ Kit v3.1 (10X Genomics), which was done following the manufacturer’s instructions. Between 13 and 15 cDNA amplification cycles were performed, and the amount of amplified cDNA used for library prep varied between 64 and 79 ng. 11 cycles of sample index PCR were performed. Libraries 1-3 (wild type) were sequenced on a NovaSeq 6000 (Illumina) and the mutant library was sequenced on a NextSeq 500 (Illumina). The sequencing configuration was 28 bp read 1 + 91 bp read 2 + 8 bp index for library 1 (wild type), which was single-indexed, and 28 bp read 1 + 90 bp read 2 + 10 bp index 1 + 10 bp index 2 for libraries 2-4 (wild type, wild type, and mutant), which were double-indexed. Targeted sequencing depth was greater than 20 thousand reads per cell.

### Quantification and statistical analysis

#### Receptor gene annotation

Zhou et al. (*56*) curated a set of gene annotations for chemoreceptors in *H. saltator*. The genes were classified into *Or*s, *Ir*s, and *Gr*s and given numbers (e.g. *HsOr322*). However, these annotations were generated using an older *H. saltator* genome assembly (Genbank accession GCA_000147195.1) (*57*), which has been since superseded by a more contiguous assembly (Genbank accession GCA_003227715.2) (*58*). Given that Zhou et al. annotations have already been used in multiple studies on ant *Or*s (*59-62*), we sought to transfer these annotations onto the new assembly and manually curate them while keeping the naming system as consistent as possible. First, we identified potential *Or* and *Gr* genes in the most up-to-date set of *H. saltator* gene annotations generated for the new assembly, HSAL51 (GEO accession GSE172309) (*63*). To do this, we scanned HSAL51 translations against the Hidden Markov Model profiles of the PFAM domains 7tm_6 (PF02949) and 7tm_7 (PF08395) (*64*) using HMMER v3.3.2 (http://hmmer.org). Then, we extracted the mRNA sequences of the candidate genes identified in this manner and performed reciprocal BLASTn search between them and the mRNA sequences from Zhou et al. using blast_rbh (*65, 66*). This strategy allowed us to assign IDs to the majority of genes, but a sizable fraction of genes remained unmatched. Also, IR genes from the new assembly were not included into the analysis until this point. Therefore, we performed manual matching and curation of the remaining genes using additional manual reciprocal BLASTn searches and tBLASTn searches against the new assembly. We also made use of the fact that most chemoreceptor genes in *H. saltator* are clustered in the genome, some clusters comprising tens of tandemly arranged genes (*56*). As an example, Zhou et al. annotations contain genes *HsOr80*, *HsOr81*, and *HsOr82*, located head-to-tail next to each other, and HSAL51 annotations contain genes *LOC105191307*, *LOC105191308*, and *LOC105191309* in the same orientation (Fig. S5B). Only *LOC105191307* and *LOC105191309* showed up as the best reciprocal BLAST hits of *HsOr80* and *HsOr82*, respectively. *HsOr81*, the gene in the middle, displayed the highest BLAST score against *LOC105191309 = HsOr82*, but it actually had higher sequence identity to *LOC105191308*. Most likely, *LOC105191308* was not the highest scoring hit because the gene model is truncated, resulting in a shorter alignment and thus a lower score. Nevertheless, we interpreted its higher sequence identity to *HsOr81* and its location relative to its well-matched neighbors as evidence of *LOC105191308* being *HsOr81*. Moreover, we used the chance to manually correct the gene model of *LOC105191308* and make it more consistent with the transcriptomic data generated earlier (*56*) and in this study. Thus, using synteny as a guide and occasionally supplementing it with additional BLAST searches, we identified the absolute majority of Zhou et al. genes in the new genome assembly, creating or updating gene models were necessary. There were several cases left when no one-to-one correspondence could be established. One example is shown in Fig. S5C: the region of the old assembly containing *HsOr142* and *HsOr143* (scaffold305:48831-53859) appears to be either an erroneous duplication in the old assembly or collapsed with a similar neighboring region in the new one. In either case, these genes are not present as distinct sequences in the new assembly. On the other hand, *HsOr181*, *HsOr182*, *HsOr183*, and *HsOr184* all had identifiable best reciprocal BLAST hits in HSAL51, but the corresponding genomic region in the new assembly contained one extra OR gene. We arbitrarily named it *HsOr182.2*, while the original best hit of *HsOr182* was named *HsOr182.1*. The final set of gene annotations was designated HSAL60 and deposited at GEO. The list of chemoreceptor genes identified in the new genome assembly is provided in Table S1 along with the description of what type of evidence was used to assign a Zhou et al.-style ID to each gene.

#### Bulk RNA-seq of pupal antennae at different developmental stages

Reads were mapped to the genome and reads overlapping gene predictions were counted using STAR v2.6.1d (*67*) with the following parameters: --alignIntronMax 7000 --quantMode GeneCounts. Since one of the sequencing batches had longer reads, reads in this batch were trimmed by appropriately specifying the --clip3pNbases parameter of STAR. Gene counts were normalized using DESeq2 v1.34.0 (*68*). To plot the temporal trend of *Or* expression in wild type vs mutant, the expression of each *Or* gene was first averaged across replicates, and then scaled to its maximum value across genotypes and stages. Next, we calculated the mean and standard deviation of the scaled expression across genes for each genotype and stage. To perform principal component analysis (PCA), we first transformed the data using the DESeq2 function vst with default parameters. Then, PCA was done either with *Or* gene set only or with the entire transcriptome. As whole transcriptome-based principal components (PCs) 1 and 2 have separated the samples by developmental stage, we identified the genes driving this separation by examining 750 genes with the largest positive loading on PC1 (presumable early stage-biased genes), 750 genes with the largest negative loading on PC2 (presumable intermediate stage-biased genes), and 750 genes with the largest negative loading on PC1 (presumable late stage-biased genes). To verify the temporal expression pattern of each of these three gene sets, we plotted the mean +/- standard deviation of their expression at each stage and for each genotype as described for the *Or* genes above. Finally, we performed GO-term enrichment using topGO v2.46.0 (*69*) in conjunction with ViSEAGO v1.8.0 (*70*). In all enrichment analyses, genes with detectable expression, defined as genes with an average sample count of at least 10, were used as the background gene set (*71*). For GO-term annotations, previously published Biological Process annotations of *H. saltator* genes (*55*) were intersected with terms present in GO.db v3.14.0 (*72*). topGOdata objects were created with the following parameters: ont=’BP’, nodeSize=5. GO-term enrichment test was performed for “early”, “intermediate”, and “late” genes separately using the following parameters: algorithm=’elim’, statistic=’Fisher’. Next, GO-terms significantly enriched in each gene set were merged using merge_enrich_terms and summarized by calculating semantic similarity using build_GO_SS and compute_SS_distances with distance argument set to ’Wang’. The merged list of GO-terms is provided in Table S2. The p-value heatmap combined with the clustered GO-term tree in Fig. 2C was generated using GOterms_heatmap with the following parameters: showIC=FALSE, showGOlabels=FALSE. To annotate the GO-term clusters, we ran compute_SS_distances on the output of GOterms_heatmap with distance argument set to “BMA”.

#### snRNA-seq pre-processing and clustering

Conversion of raw sequencing data to FASTQ, creation of the transcriptome index, and read counting to generate expression matrix was done in cellranger v7.0.0 with default parameters (*73*). Subsequent analyses were done using scanpy v1.8.2 (*74*). First, cell-wise and gene-wise filtering was applied as follows. Cells with fewer than 750 detected genes, greater than 12,500 UMIs, or greater than 2.5% mitochondrial reads were removed from the analysis. Then, genes annotated in the NCBI file All_Invertebrates.gene_info (https://ftp.ncbi.nih.gov/gene/DATA/GENE_INFO/Invertebrates/All_Invertebrates.gene_info.gz) as “large subunit ribosomal RNA” were removed. First, the different libraries were merged (AnnData.concatenate) and analyzed without performing any batch correction, except what scanpy documentation calls “lightweight batch correction” at the stage of variable gene selection. The combined counts were depth-normalized (CP10k) and log+1 transformed using default parameters. Highly variable gene selection was done with the following parameters: layer=’counts’, batch_key=’orco’, flavor=’seurat_v3’, n_top_genes=2000, where ‘orco’ is the metadata attribute that encodes whether the library is wild type or mutant. PCA was performed, and the optimal number of PCs (12) was chosen as the point at which the proportion of variance explained by each PC plateaued. The neighborhood graph and UMAP were computed with default parameters except the number of PCs. Plotting library ID and quality control metrics on the UMAP revealed the following: 1) even wild type libraries prepared from different batches of biological material on different days, i.e. library 1 vs libraries 2 and 3, exhibited pronounced batch effects; 2) quality metrics, e.g. the number of UMIs, appeared to have a strong effect on cell clustering. Thus, we reanalyzed the data by applying batch correction and increasing the stringency of cell-wise filtering. In addition to the filtering cutoffs applied above, cells with greater than 2,750 detected genes, greater than 9,500 UMIs, or greater than 1.25% mitochondrial reads were removed. The filtered raw counts were depth-normalized (CP10k) and log+1 transformed using default parameters. Highly variable gene selection was done with the following parameters: layer=’counts’, batch_key=’sample’, flavor=’seurat_v3’, n_top_genes=2000, where ‘sample’ is the metadata attribute that contains the library ID (1, 2, 3, or 4). Then, scvi-tools v0.16.2 (*75*) was used to set up an scVI model with layer=’counts’, batch_key=’sample’, and then train the model with the following parameters: max_epochs=800, early_stopping=True, deterministic=True. Obtained latent representation was used to compute the neighborhood graph and UMAP, and Leiden clustering was performed with an arbitrary selected resolution of 5.

#### Cell type annotation

Next, we set out to annotate the cell types. First, cells were broadly split into neurons and non-neurons. Neurons were defined as cells that belonged to clusters that simultaneously expressed *LOC105189534 / nSyb*, *LOC105190174 / fne*, *LOC105183410 / Syt1*, and *LOC105183587 / onecut*, while the rest of the cells were classified as non-neuronal cells. Next, given that different ORN types may primarily differ by the expression of only one or several receptor genes and that they may not be represented in our data set in large numbers, we decided against assigning ORN types to clusters of cells and instead classified each individual cell. To overcome the sparsity of 10X data, we employed the following strategy: a Mann-Whitney U test (scipy.stats.mannwhitneyu) was performed for each receptor gene to compare its expression in the previously defined neighborhood of the focal cell and in a randomly chosen set of 100 wild-type non-neuronal cells. A p-value cutoff of 0.05 was used to label each cell as expressing or not expressing a given gene (Fig. S2A). Then, each individual neuron was classified in the following way, where + stand for expression (neighborhood p-value < 0.05) and - stands for the lack of expression:

| ***LOC105188823 / Rh50*** | ***LOC105188598 / nompC*** | **At least one GR** | **At least one Irco** | **At least one tuning IR** | **Orco** | **At least one tuning OR** | **Cell type assigned** |
| --- | --- | --- | --- | --- | --- | --- | --- |
| + | + or - | + or - | + or - | + or - | + or - | + or - | ammonia-sensory neurons |
| - | + | + or - | + or - | + or - | + or - | + or - | mechanosensory neurons |
| - | - | - | - | - | + | + | chemosensory neurons (OrX+, Orco+) |
| - | - | + | - | - | + | + | chemosensory neurons (OrX+, Orco+, GrX+) |
| - | - | - | + | - | + | + | chemosensory neurons (OrX+, Orco+, Irco+) |
| - | - | - | + | + | + | - | chemosensory neurons (IrX+, Irco+, Orco+) |
| - | - | + | + | + | - | - | chemosensory neurons (IrX+, Irco+, GrX+) |
| - | - | + | + | + | - | + | chemosensory neurons (IrX+, Irco+, GrX+, OrX+) |
| - | - | + | + | - | - | - | chemosensory neurons (GrX+, Irco+) |
| - | - | + | + | - | - | + | chemosensory neurons (GrX+, Irco+, OrX+) |
| - | - | all other patterns |  |  |  |  | unclassified |

*LOC105188823 / Rh50* and *LOC105188598 / nompC* were chosen as markers of ammonia- and mechanosensory neurons, respectively, following (*52, 76*). Finally, cells that belong to clusters expressing *LOC105181500 / repo* were classified as glia (*77*), *LOC105191850 / Mhc* was used as the marker of muscle cells (*32*), *LOC105181616 / grh*-positive cells were classified as epithelium (*32*), and the expression of *LOC105187024 / sv* was used to identify neuronal support cells (*32, 42*).

#### OR phylogeny

To reconstruct the phylogeny of the HSAL60 *Or* genes, we first identified the longest predicted transcript of each gene. Then, we translated them using TransDecoder v5.5.0 (https://github.com/TransDecoder/TransDecoder) and only retained a single translation per transcript by passing the --single_best_only flag to TransDecoder.Predict. The translations were aligned using MAFFT v7.508 (*78*) with default parameters. The alignment was visually examined using Jalview v2.11.2.4 (*79*) and sites deemed non-informative were manually removed. Then, phylogenetic tree was built by running RAxML v8.2.12 (*80*) with the following parameters: -f a, -m PROTGAMMAAUTO, -# 100. The best-scoring tree was re-rooted with Orco as an outgroup using FigTree v1.4.4 (http://tree.bio.ed.ac.uk/software/figtree/). The tree in the Newick format is provided as Data S1.

#### Bulk RNA-seq of adult antennae

Reads were mapped to the genome and reads overlapping gene predictions were counted using STAR v2.6.1d (*67*) with the following parameters: -- alignIntronMax 7000 --quantMode GeneCounts. The counts were normalized using DESeq2 v1.34.0 (*68*) and overlaid on the OR tree using ggtree v3.2.1 (*81*).

#### Identification of Orco-independent ORs

We used both the single-nucleus data and the bulk data to identify the Orco-independent genes. Raw single-nucleus counts were depth-normalized (CP10k) and the average expression of each OR gene in mutant cells was divided by its average expression in wild-type cells. The resulting fold change values were log2-transformed. Only genes with non-zero expression in both wild-type and mutant samples were considered. Similarly, FPKM values from the bulk data (see above) were averaged across wild-type and mutant samples, and log2 fold change between mutant and wild type was calculated for each OR gene. Plotting log2 fold changes in single-nucleus and bulk data revealed a highly significant positive relationship (Pearson R^2^ = 0.21, p < 10^-13^, Spearman R^2^ = 0.17, p < 10^-11^) (Fig. S3A). To select genes that exhibited the smallest amount of change in the mutant, we drew an arbitrary cutoff for both data sets, thus identifying genes that are either downregulated the least or upregulated in the mutant (Fig. S3A). Such genes were designated as Orco-independent. To identify the markers of non-neuronal cells that express *HsOr370*, we performed iterative marker searches using sc.tl.rank_genes_groups with either ‘rest’ or various individual clusters as the reference. The expression specificity was visually assessed by plotting the expression of potential marker genes on the UMAP.

## KEY RESOURCE TABLE

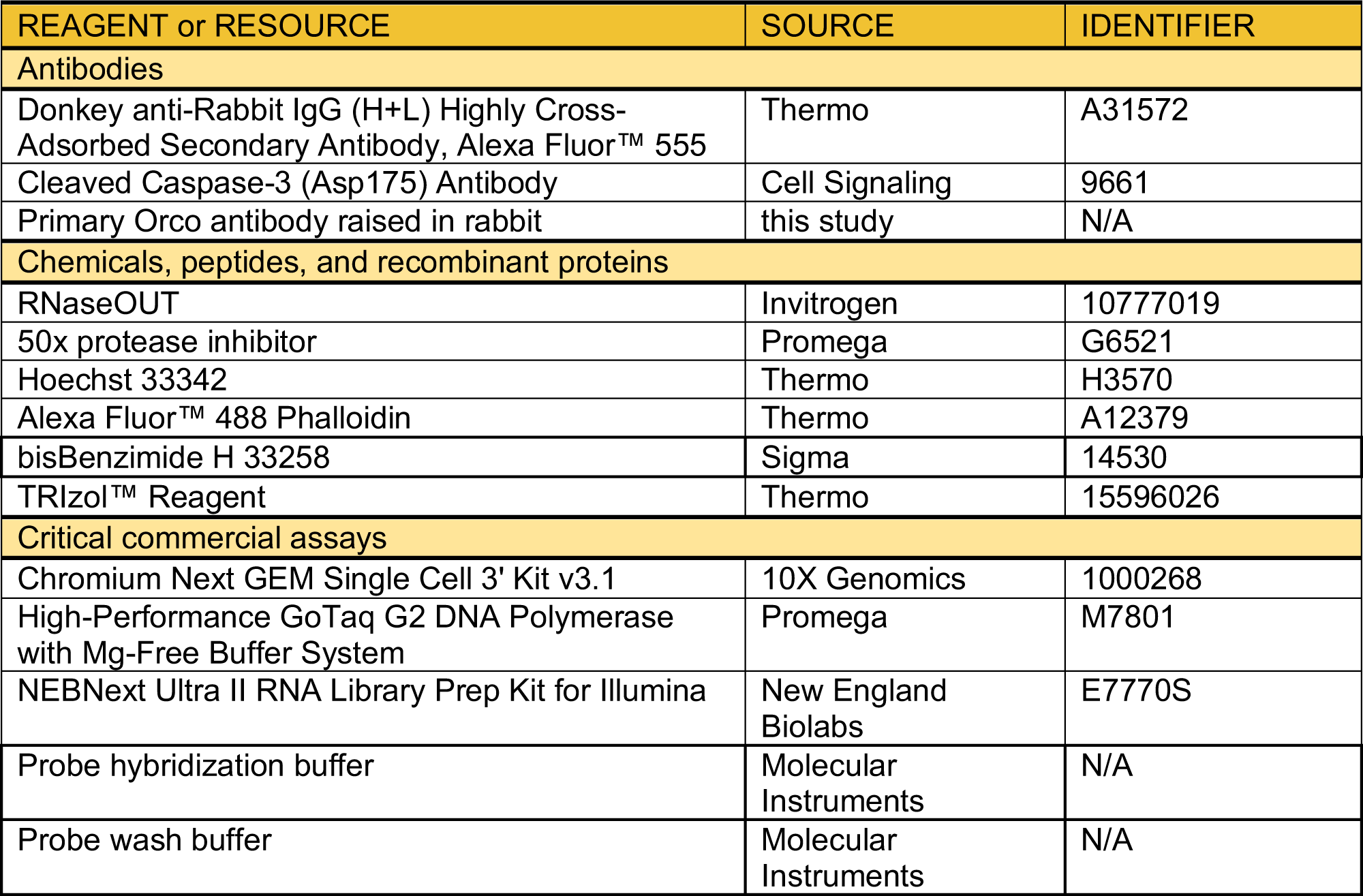
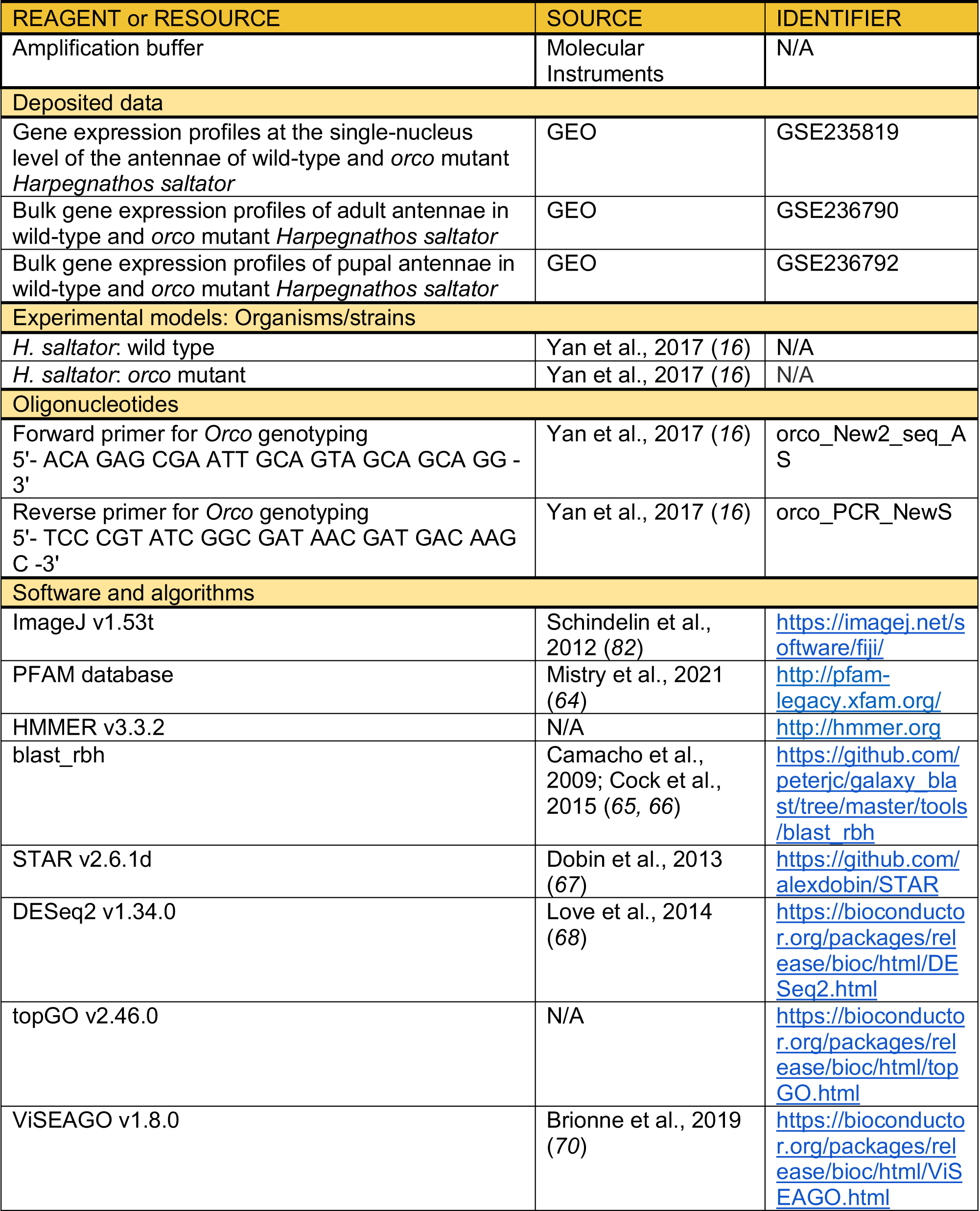
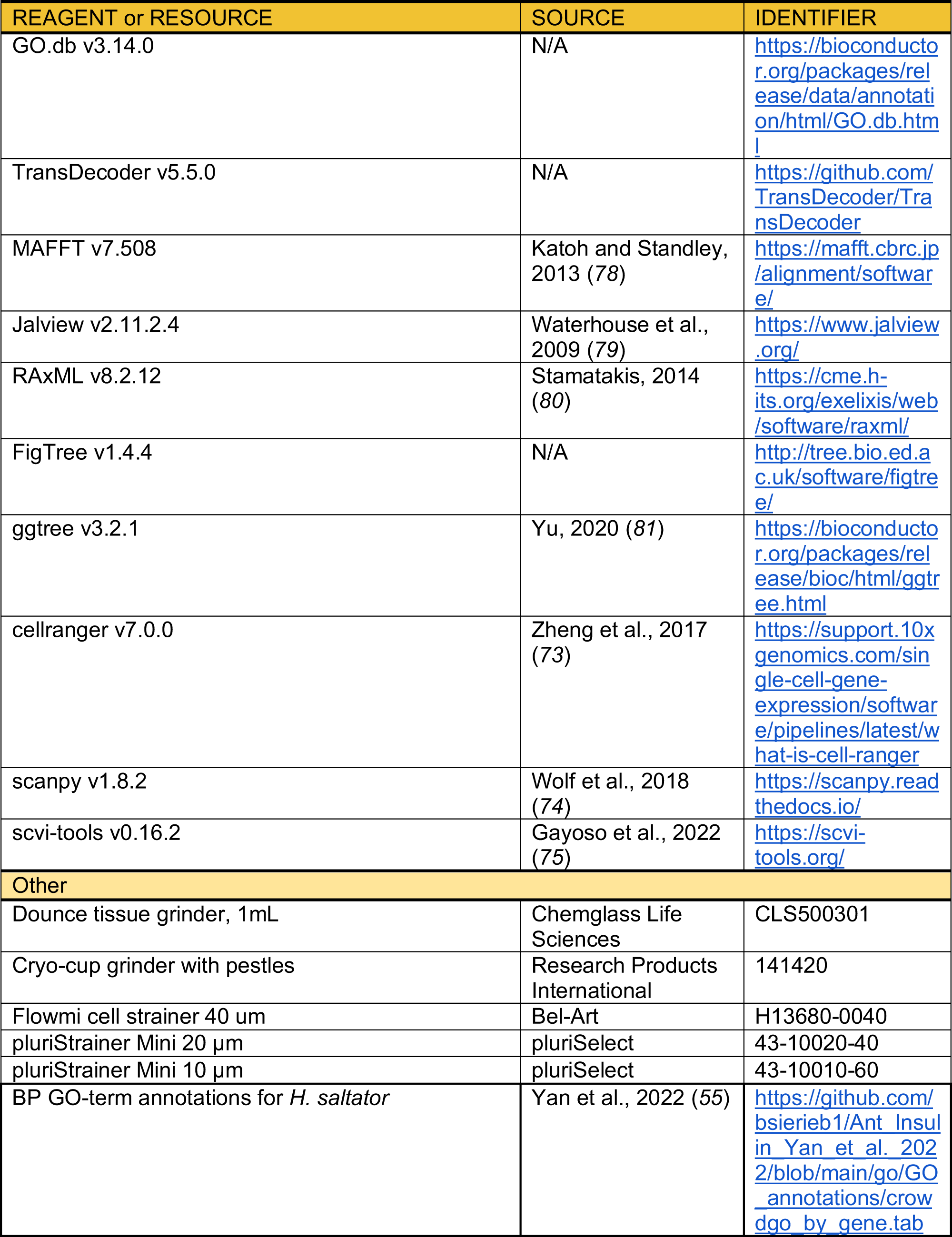

## ACKNOWLEDGMENTS

This study was funded by NIH grant R01DC020203 and NSF I/UCRC CAMTech grant IIP1821914 to H.Y. B.S. was supported by a Long-Term Fellowship LT000010/2020-L from the Human Frontier Science Program. K.S. was supported by a Predoctoral Training Program T32DC015994 from NIH. We thank Claude Desplan and his laboratory at NYU for helpful discussions and technical support.

## AUTHOR CONTRIBUTIONS

Conceptualization: BS, KRS, HY; Methodology: BS, KRS, HY; Investigation: BS, KRS, OK, JM, SJ, HY; Visualization: BS, KRS, OK; Supervision: HY; Writing: BS, KRS, HY

## SUPPLEMENTARY MATERIALS

**Figure S1.**
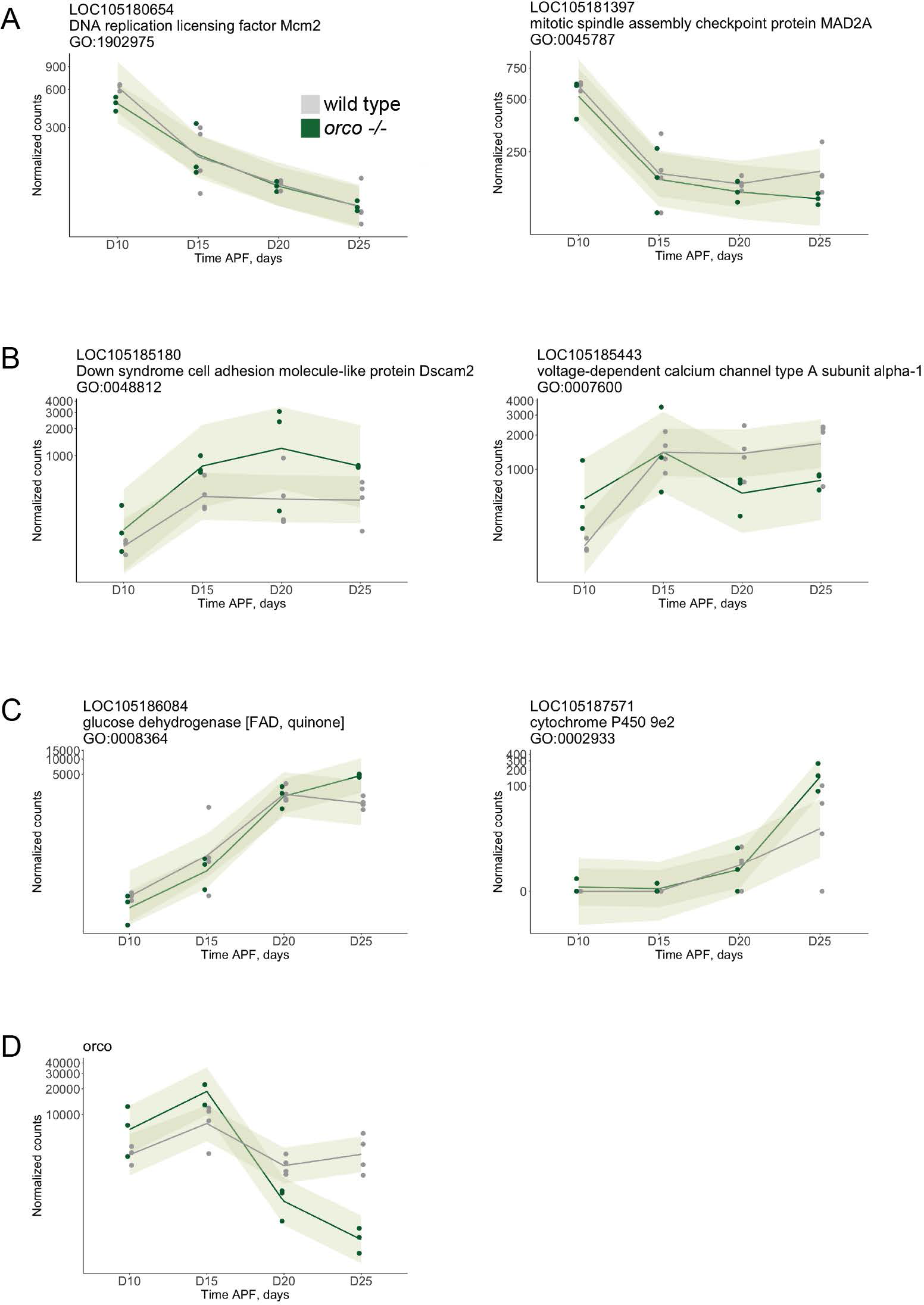
Gene expression in wild type and *orco* mutant antennae at different time points. **(A-C)** Expression patterns of selected genes from “early” (A), “intermediate” (B), and “late” (C) gene sets as defined in Fig. 2B. The title of each plot contains gene ID, NCBI annotation, and an associated GO-term significantly enriched in the corresponding gene set. Note that genes shown may be associated with additional GO-terms.

**(D)** Expression pattern of *Orco*.

The Y axis in all panels displays DESeq2-normalized gene counts. Each point corresponds to a sample. The lines are LOESS regression lines for the wild type and the mutant, respectively, and the shaded areas depict 95% confidence intervals.

**Figure S2.**
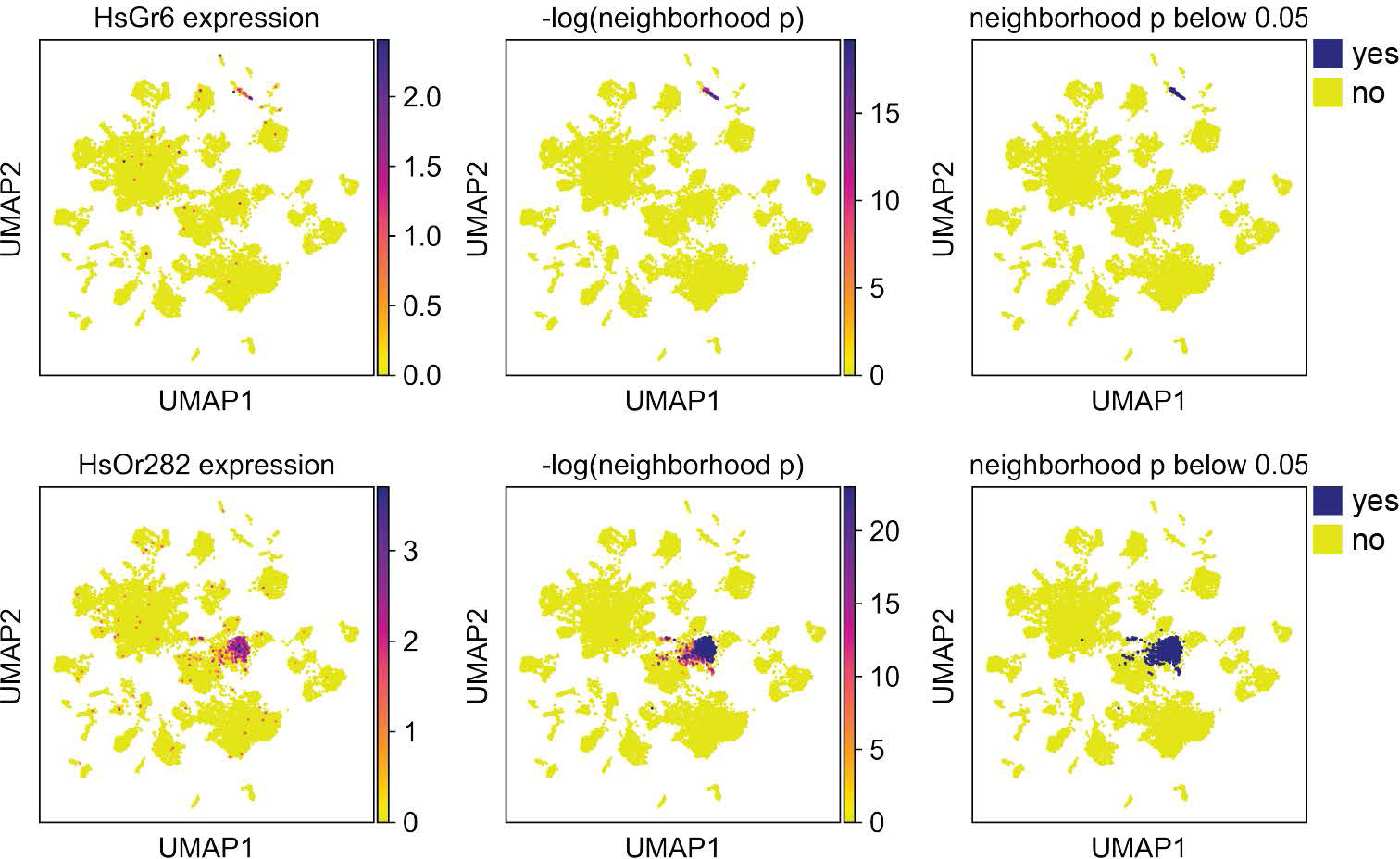
Binarization of receptor gene expression. UMAP plots showing the expression of two selected receptor genes. Plots in the left column show depth-normalized and log-transformed counts. Plots in the middle column show the negative logarithm of the p-value obtained from the Mann-Whitney U test between the neighborhood of each cell and a randomly chosen set of 100 wild-type non-neuronal cells (see Methods for details). Plots in the right column show binarized p-values from the middle column (above or below 0.05), which were used to classify cells as expressing or not expressing different receptor classes shown in fig. 3C.

**Figure S3.**
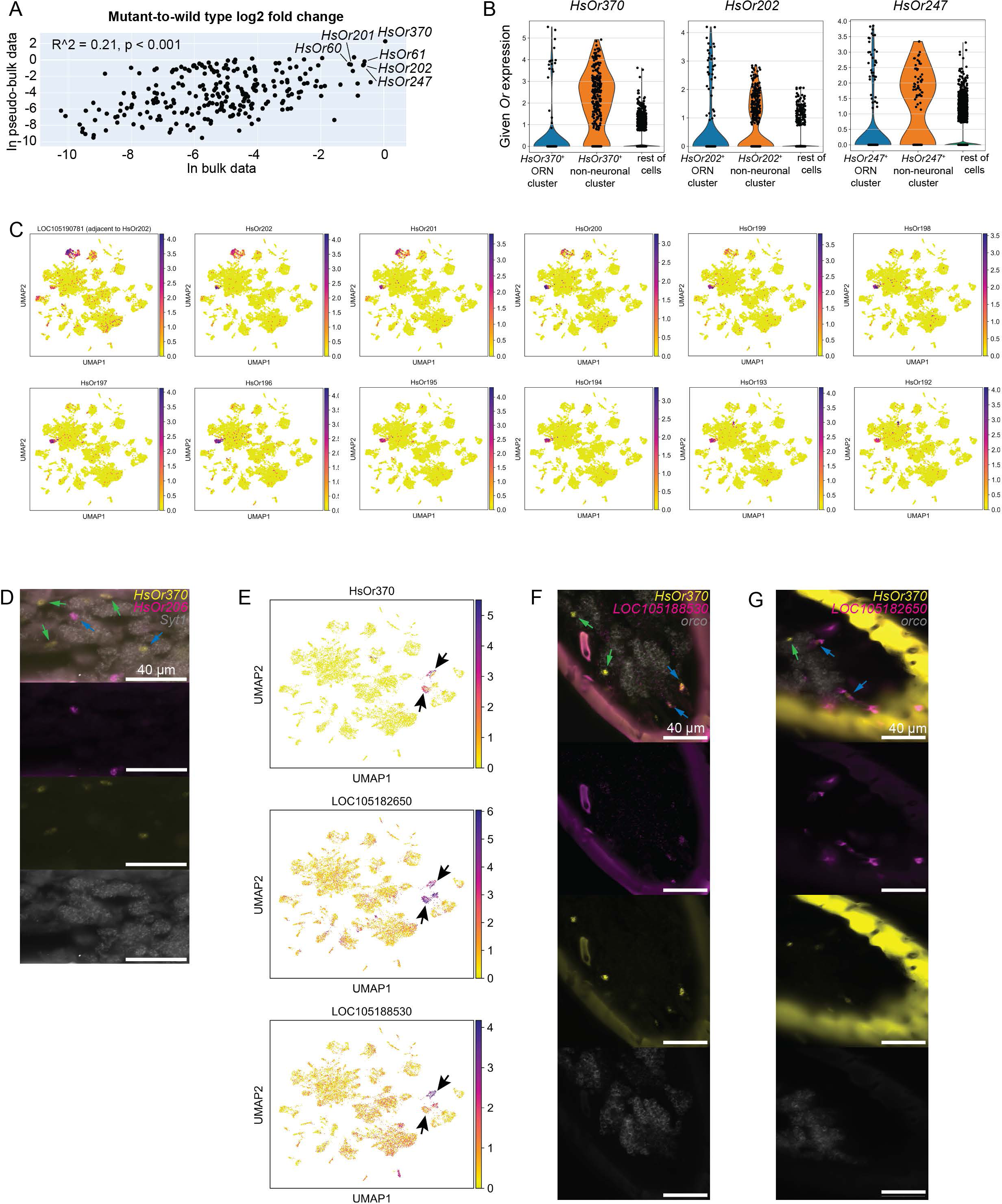
Non-neuronal expression of chemoreceptor genes in *H. saltator*. **(A)** Correlation between “Orco-independence” (expressed as a log2 fold change in *Or* expression between mutant and WT) in bulk data and “Orco-independence” in pseudo-bulk data. **(B)** Expression level of selected “Orco-independent” *Or*s in ORNs and non-neurons. **(C)** UMAP plots showing the expression of *Or*s *HsOr202-HsOr193* and their upstream non-*Or* neighbor. **(D)** Representative HCR RNA-FISH images of sectioned WT adult antennae illustrating the relationships between an Orco-dependent gene (*HsOr206*, magenta), an Orco-independent gene (*HsOr370*, yellow), and a neuronal marker (*Syt1*, grey). The Orco-dependent gene always co-localizes with the neuronal marker (left blue arrow). The Orco-independent gene may co-localize with the neuronal marker (right blue arrow) or without (green arrows). **(E)** UMAP plots showing the expression of *HsOr370* and of two genes marking the *HsOr370*-positive support cells. Arrows point at the support cell clusters that express *HsOr370*. **(F-G)** Representative HCR RNA-FISH images of sectioned WT adult antennae. Here, we show the relationship between an Orco-independent gene (*HsOr370*, yellow), identified support cell markers (magenta) and a neuronal marker (orco, grey). Green arrows show cases of *HsOr370* expression in neuronal cells. Blue arrows demonstrate cases where *HsOr370* co-localizes (D) or “nests” (E) with support cell markers, but not the neuronal marker.

**Figure S4.**
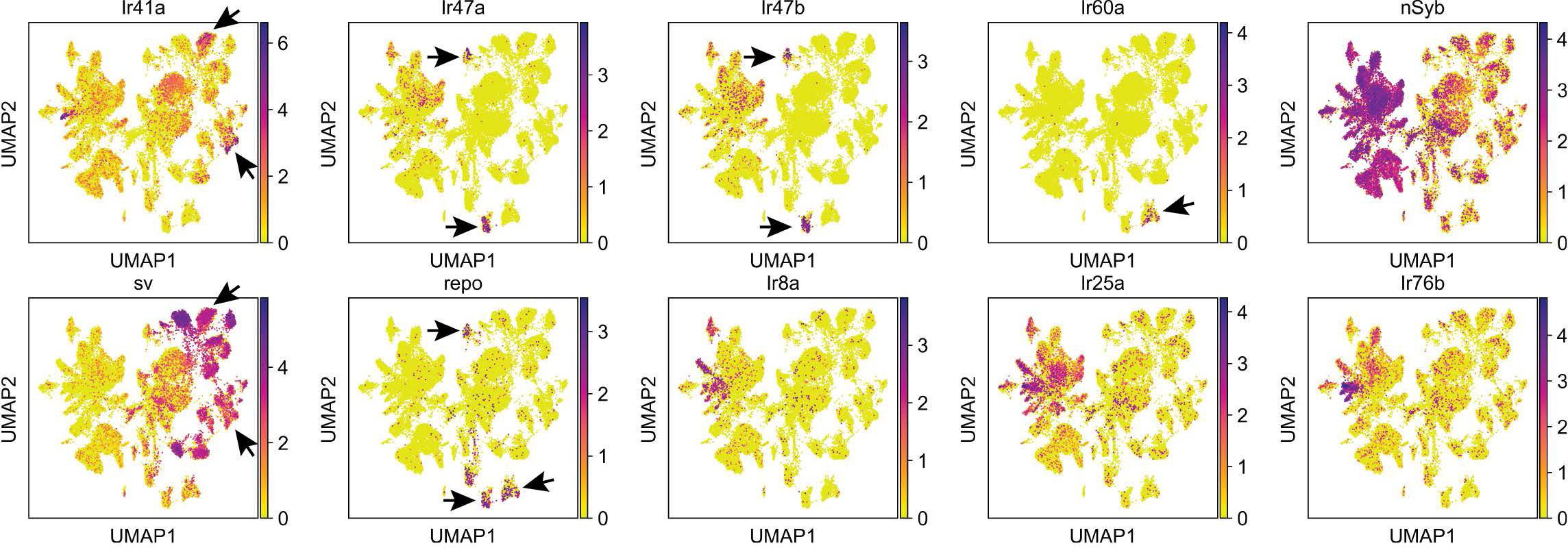
Non-neuronal expression in *Drosophila melanogaster*. UMAP plots showing the expression of the four non-neuronally expressed *Ir*s, a neuronal marker *nSyb*, a support cell marker *sv*, a glial cell marker *repo*, and all three *irco*s in *Drosophila*.

**Figure S5.**
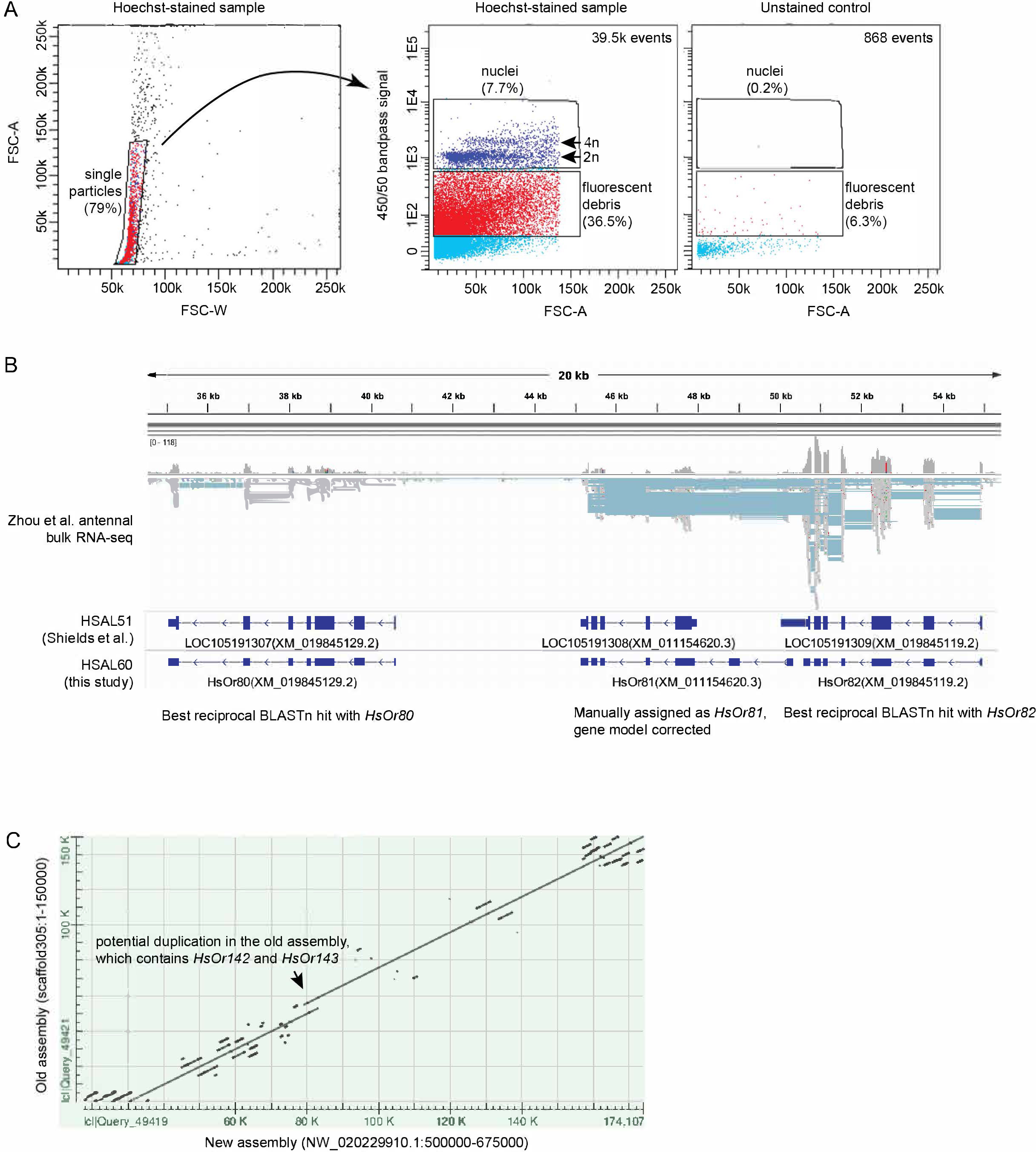
Additional experimental details. **(A)** FACS gating. FSC = forward scatter, A = area, W = width, 2n = putative diploid nuclei, 4n = putative tetraploid nuclei. The “nuclei” gate was used for sorting. **(B)** Genome browser snapshot showing the locus that contains *HsOr80, HsOr81,* and *HsOr82* along with aligned bulk RNA-seq data, HSAL51 and HSAL60 gene annotations, and the description of evidence used to transfer gene IDs from Zhou et al. onto the HSAL60 annotations. **(C)** Dot plot output of BLAST between the region that contains *HsOr142* and *HsOr143* in the old genome assembly and the homologous region in the new assembly.

**Table S1. List of *Gr*, *Ir*, and *Or* genes in the HSAL60 annotation set.**

**Table S2. List of GO-terms enriched in “early”, “intermediate”, and “late” gene sets.**

**Data S1. Phylogenetic tree of *Or* genes in the Newick format.**

